# Sugar intake elicits a small-scale search behavior in flies and honey bees that involves capabilities found in large-scale navigation

**DOI:** 10.1101/171215

**Authors:** Axel Brockmann, Satoshi Murata, Naomi Murashima, Ravi Kumar Boyapati, Manal Shakeel, Nikhil G. Prabhu, Jacob J. Herman, Pallab Basu, Teiichi Tanimura

## Abstract

Social insects, particularly bees and ants, show exceptional large-scale navigational skills to find and carry back food to their nests. Honey bees further evolved a symbolic communication to direct nest mates to attractive food sources. Till now it is unclear how these capabilities evolved. Sixty years ago, Vincent Dethier demonstrated that a small-scale sugar-elicited search behavior identified in flies shows remarkable similarities with honey bee dance behavior. Those findings suggested that both behaviors are based on common mechanisms and are likely evolutionary related. We now present for the first time a detailed comparison of the sugar-elicited search behavior in *Drosophila melanogaster* and *Apis mellifera*. In both species, intake of sugar elicits a complex of searching responses. The most obvious response was an increase in turning frequency, but more importantly we found that flies and honey bees returned to the location of the sugar drop. They even returned to the food location when we prevented them from using visual and chemosensory cues indicating that this small scale local search involves path integration mechanisms. Finally, we show that visual landmarks presented in the vicinity of the sugar drop affected the search trajectory and in honey bees the sugar intake induced learning of landmarks. Together, our experiments indicate that the sugar-elicited local search exhibits two major behavioral capabilities of large-scale navigation, path integration and landmark orientation.

**Significance Statement:** To search for food social insects evolved sophisticated strategies of spatial orientation and large-scale navigation. We now show that even a small-scale local search behavior in solitary flies and social honey bees involves path integration and landmark learning two major mechanisms of large-scale navigation. We propose that in the future sugar-elicited local search can be used to identify neural circuits involved in navigation, path integration, and landmark learning.

## Introduction

Food search behaviors are the most successful experimental paradigms to study navigation and spatial memory in insects and vertebrates (1,2). Search for food can be separated into two distinct phases: a hunger induced large-scale search for food sources and a food intake elicited local search for more food (3,4). Interestingly, sixty years ago the American entomologist Vincent Dethier suggested that a simple sugar-elicited local search behavior observed in solitary flies might represent an ancestral behavioral locomotor pattern that was co-opted during evolution into the honey bee dance behavior which communicates the distance and direction to a food source (4,5). Sugar-elicited search and honey bee dance are similar in that they are initiated after the intake of food and include a stereotypic turning behavior; which is modulated by the reward value of the food source and the internal state of the individual (4-6).

We got interested in three questions, first do honey bees actually show sugar-elicited search behavior, second if so, how similar is this search behavior in solitary flies and social honey bees, and third is the search behavior based on a simple increase in turning frequency or does it involve more complex mechanisms of spatial orientation. To answer these questions we first developed similar behavioral assays for flies and bees and then tested different aspects of the search behavior: (a) effect of sugar concentration on the intensity of search behavior, (b) effect of lightning condition on search trajectories, (c) which sensory systems are necessary for guiding the search trajectory, and (d) does the sugar intake has the capability to induce learning processes that might affect the following search trajectory?

Our experiments show that social honey bees initiated a search behavior after ingesting a drop of sugar which is very similar to that of solitary flies. Small differences in this search behavior are likely due to differences in more general aspects of locomotor behavior and nutritional physiology. More importantly, our analyses indicate that sugar-elicited search behavior is not just a simple turning behavior but involves a set of complementary responses: change in turning frequency, initiation of path integration, and initiation of landmark learning. These findings suggest that this small-scale spatial orientation involves two major behavioral capabilities and strategies involved in large-scale navigation (1,5,7,8). Thus, sugar-elicited search behavior promises to be a very fruitful behavioral paradigm to identify neural and molecular mechanisms involved in small- and large-scale navigation and possibly also honey bee dance communication.

## Results

### Sugar-elicited search behavior in flies and honey bees

Starved flies and feeder arriving (i.e. hungry) honey bee nectar foragers showed a variety of locomotor responses after the intake of a little drop of water or sugar-solution. The least response was a short relatively straight walking trail and a rapid flying off. Most intricate trajectories consisted of initially small circles around the location of the sugar drop which increased with time (Fig. 1*B* also *Fig. 2*; see movies S1-S4). Flies walked in bouts frequently stopping for a short time. These walking bouts were relatively straight. Flies mainly changed directions performing sharp turns during the stops (Fig 1*A*/*C*; 3,9,10). In all our experiments flies showed a high amount of grooming which is likely due to the relatively large sugar water volume fed (11,12). In contrast, honey bees constantly moved in a meandering way (Fig 1*A*/*C*). Search trajectories of *Apis mellifera* workers showed a more uniform distribution of speed than the trajectories of *Drosophila* (Fig. 1*D*; Kolmogorov-Smirnov test; *P* < 0.0001). Second, the distribution of z-value of curvature for the search trajectory was significant flatter in bees than flies (Fig 1*C*; Mann-Whitney U test, P < 0.0001). For both species, scatter plots of speed (z-values) vs curvature (z-values) indicate an increase in variation of curvature with speed, i.e. sharper turns were taken with higher speed. Thus, flies and bees do not slow down but speed up when making turns during search behavior, which might suggest a state of heightened arousal.

**Fig. 1.**
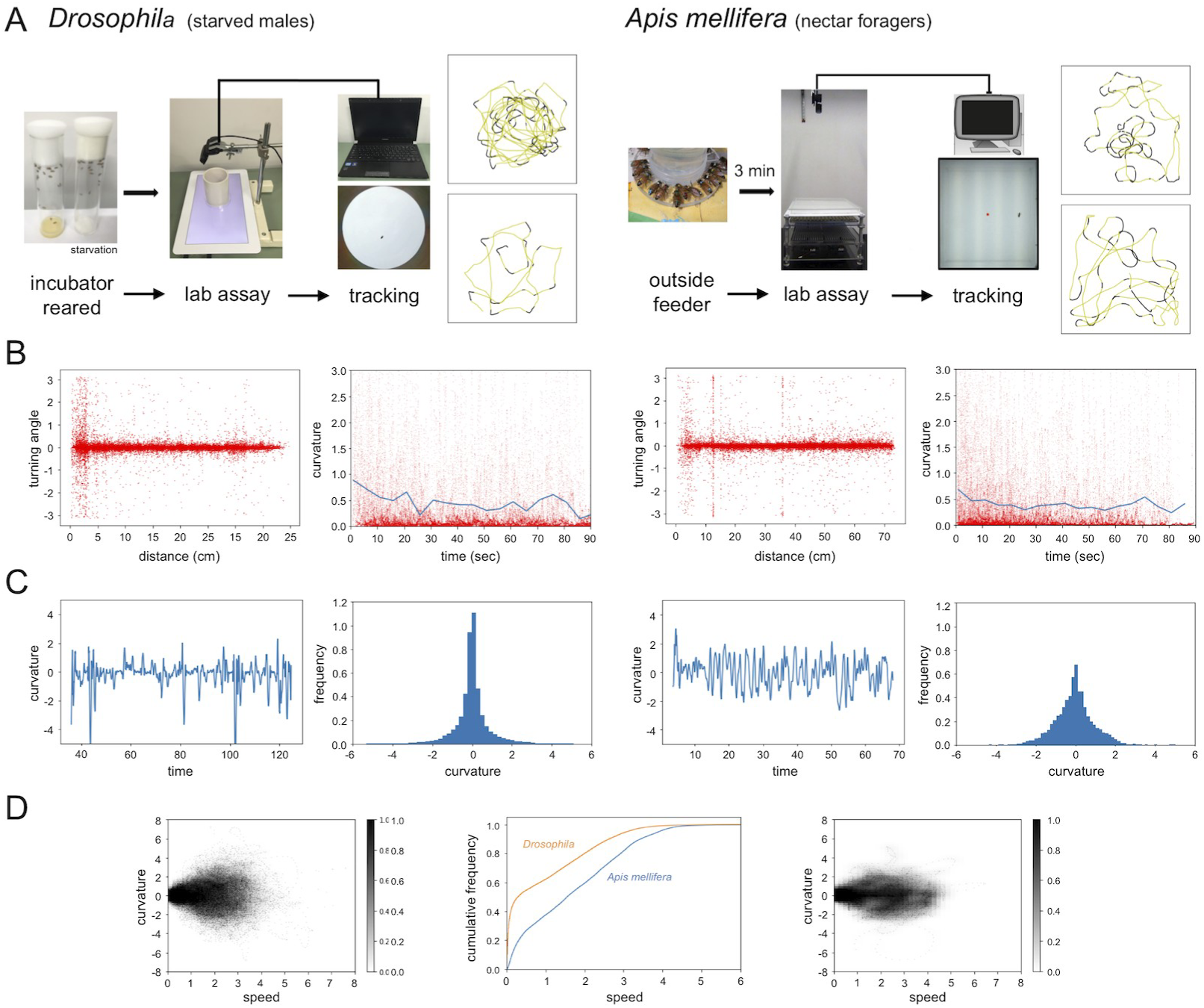
Sugar-elicited search behavior in flies and honey bees. (A) Scheme of experimental procedures and two selected SeS-walking trajectories of Drosophila males and *Apis mellifera* workers (B) Curvature (z-values) versus time plot of an individual search trajectory and histogram of curvature for the all experimental trajectories (*Drosophila n* = 20; *Apis mellifera n* = 20). Distribution of sharp turnings showed significant difference between Drosophila and *Apis mellifera* (Mann-Whitney U test, *P* < 0.0001) (C) Plots of turning angle versus radial distance from the sugar drop location and curvature versus time. Both flies and bees show a higher rate of turning close to the location of the sugar drop and both show a higher turning behavior at the beginning of the search. (D) Plot of curvature versus speed distribution and cumulative histogram of speed distribution. Apis mellifera workers show a more uniform distribution of speed than *Drosophila* (Kolmogorov-Smirnov test; *P* < 0.0001). Drosophila walking trajectories were characterized by frequent stops (i.a. bouts of grooming) and high speed turns. In contrast, *Apis mellifera* workers constantly moved.

**Fig. 2.**
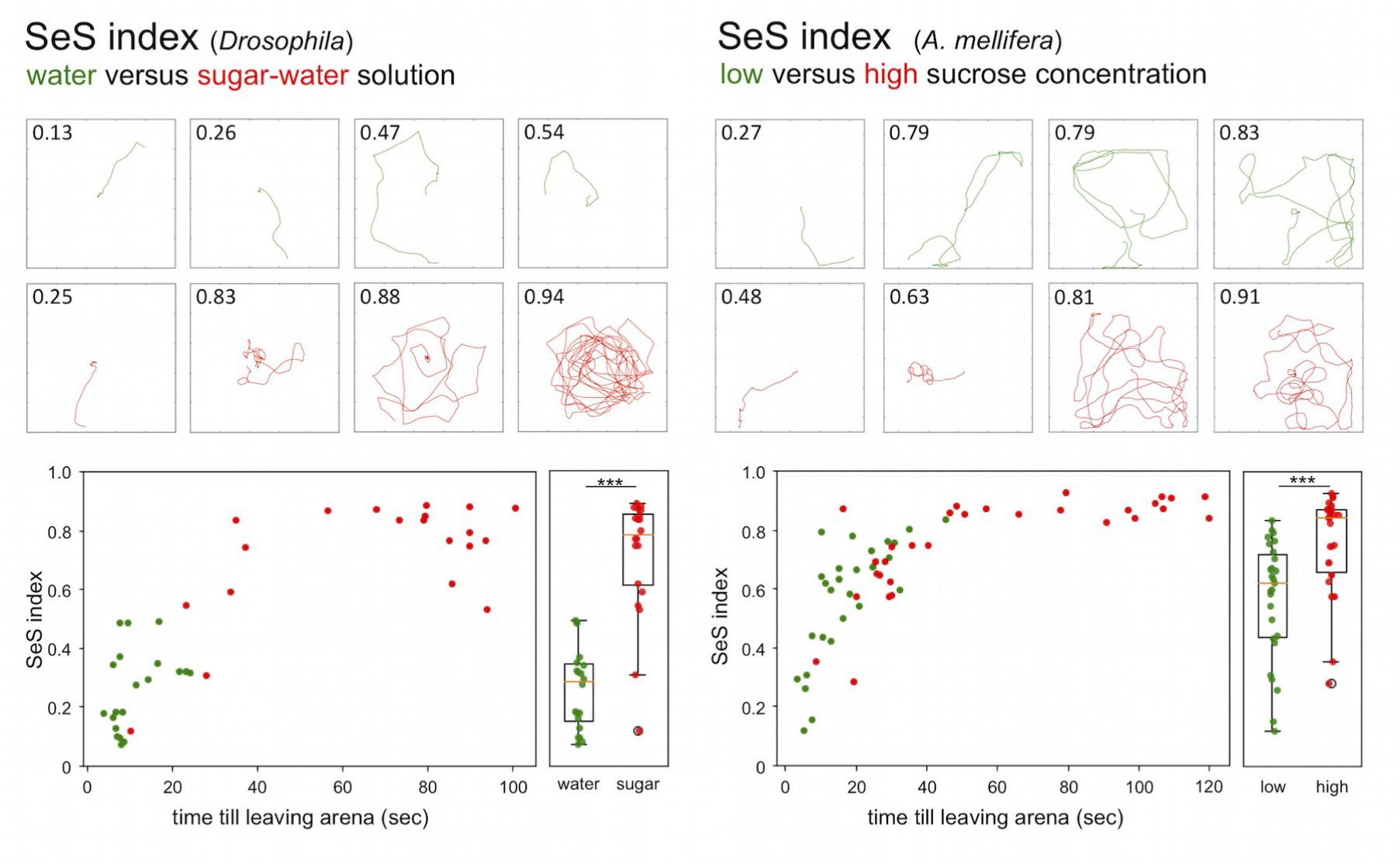
Comparison of sugar-elicited search behavior induced by different food rewards. Selected walking trajectories with SeS-indices. Higher SeS-indices indicate a more meandering walking path characteristic for a local search. Drosophila: green = water; red = sugar solution. Honey bee: green = 0.2 M; red = 2.0 M. SeS-indices clearly separate walking trajectories of flies having ingested water and sugar solution as well as walking trajectories of honey bees fed a 0.2 M or 2.0 M sugar solution. *Drosophila*: water-fed, *n* = 20, median SeS-index = 0.23; sugar-fed, *n* = 20, median SeS-index = 0.77; Mann-Whitney U test *P*_(2)_< 0.0001. *A. mellifera*: 0.2 M group, *n* = 25, median SeS-index = 0.61; 2.0 M group, *n* = 27, median SeS-index = 0.81; Mann-Whitney U test *P*_(2)_= 0.0002.

In addition to these analyses, we developed a sugar-elicited search index (SeS-index) to simplify comparison of sugar-elicited search behavior among treatment groups, behavioral phenotypes and genetic strains. The SeS-index ranges from 0 to 1 and indicates the degree of convolution of the walking trajectory. As flies and honey bees differ in their general walking pattern, we developed two slightly different formulas of SeS-index for the two species (for details see, *Materials and Methods*). In a first experiment, we compared search trajectories induced by water and sugar-water solutions. *Drosophila* males were presented either with a 0.1 ul drop of water or 0.2 M sugar-water solution. Water- and food-starved flies accepted and ingested the drop of water but left the location in a more or less straight path, and in most cases quickly started flying off. In contrast, ingestion of sugar-water initiated a complex convoluted walking path. The SeS-index was significantly different between both groups (water: *n* = 20; sugar fed, *n* = 20, Mann-Whitney U test *P*_(2)_ < 0.0001; Fig. 2). These results suggest that sugar but not water induces local search behavior in *Drosophila* (12).

*A. mellifera* nectar foragers were presented with a 3 ul drop of water, low (0.2 M) or high (2 M) concentrated sugar-water solution. Nectar foragers (*n* = 30), which we had collected at a 1M feeder, never accepted a drop of water in our experiments. This corresponds with earlier studies showing that more than 90% of foragers visiting a 50% (∼ 1.46 M) sugar-water feeder did not accept water in a laboratory assay (13). When presented with a low or high sugar water most of the foragers initiated a complex meandering trajectory with high SeS-indices. The intensity of the search was significantly higher in the high sugar concentration group (0.2 M group; *n* = 25; 2.0 M group; *n* = 27; Mann-Whitney U test *P*_(2)_ = 0.0002; Fig. 2). For both honey bee groups, we found a positive correlation between SeS-indices and stay time (= time till leaving the arena; 0.2 M; r = 0.87; Mann-Whitney U test *P* < 0.0001; 2.0 M; r = 0.89; *P* < 0.0001). In contrast, SeS-index and stay time did not correlate for the water-fed flies (r = 0.28; *P* = 0.28) and only showed a weak correlation for sugar-fed flies (r = 0.58; *P* < 0.01). This finding suggests that search intensity and staying time at a location are likely independent behavioral units and are differently regulated in solitary flies and social honey bees.

### Effects of lighting conditions on search behavior

Next, we compared search trajectories performed under light and dark (infrared) conditions. Flies and honey bees initiated search behavior under both conditions and the SeS-indices did not significantly differ among conditions for each species. However, flies stayed closer to the location of the sugar drop in the dark condition (light: *n* = 14; dark: *n* = 12; Mann-Whitney U test *P*_(2)_ = 0.018, Fig. 3*A*, see movie S5), whereas honey bees walked further away from the sugar drop (light: *n* = 9; dark: *n* = 10; Mann-Whitney U test *P*_(2)_ = 0.037, Fig. 3*A*, see movie S6).

**Fig. 3.**
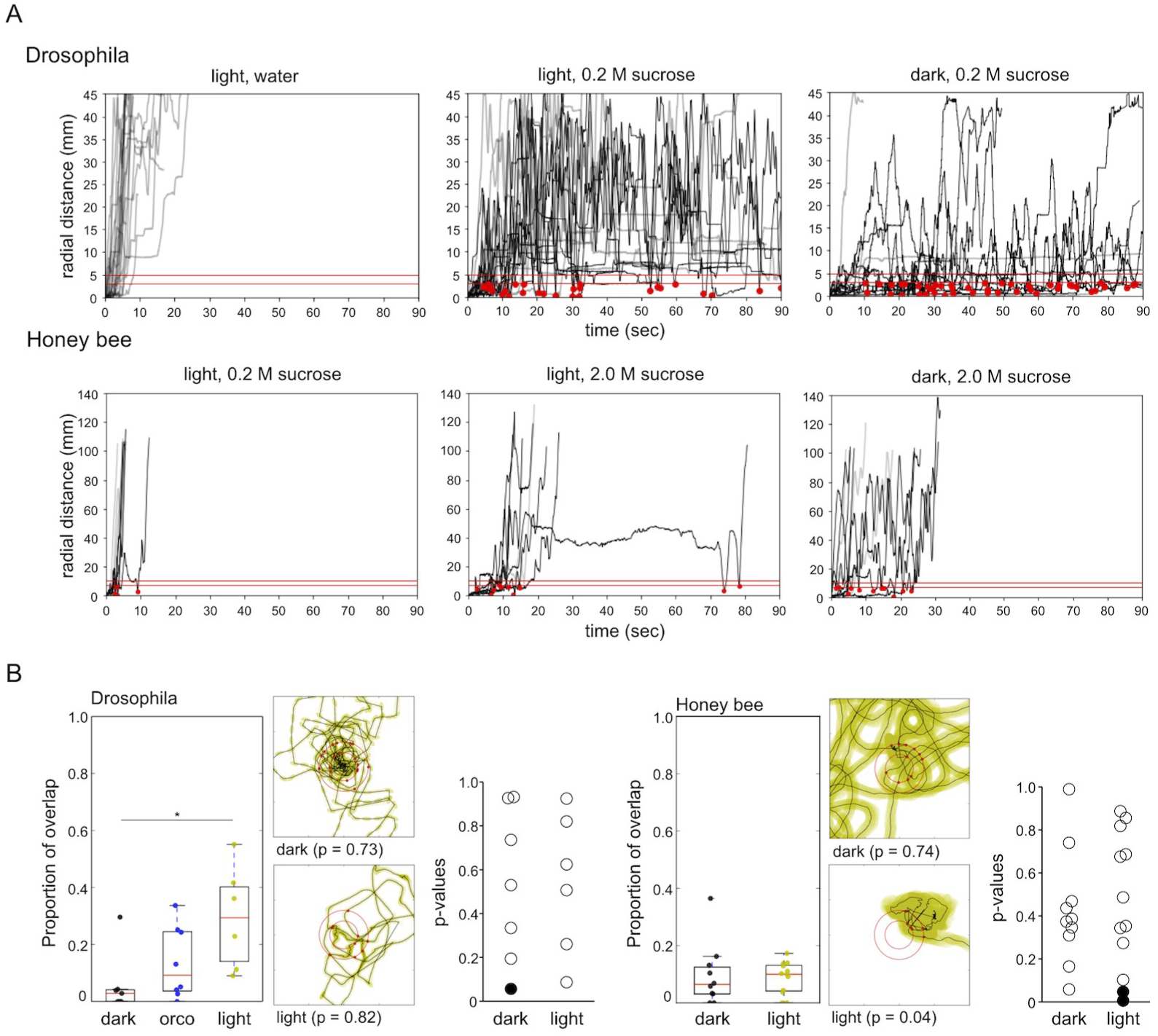
Effects of lighting conditions on search behavior. (A) Time plots of radial distances for search trajectories induced by different food rewards and different lighting conditions. Graphs comprise trajectories of flies and bees tested under the respective conditions (*Drosophila*: light water *n* = 10; light sugar *n* = 11; dark sugar *n* = 11; *A. mellifera*: light 0.2 M sugar *n* = 10; light 2.0 M sugar *n* = 9; dark 2.0 M sugar *n* = 10). Black lines: walking trajectories with returns to location of the sugar drop; grey lines: no returns. Red lines: threshold distances to identify a return to the sugar drop location (see experimental procedures). The upper red line (r1): distance (r1 *Drosophila* = 4 mm, r1 *A. mellifera* = 12 mm) the animal had to walk away from the sugar drop location before returning; lower red line (r0): distance (r0 *Drosophila* = 2.5 mm, r0 *A. mellifera* = 7 mm) the animal has to reach to be counted as a return. Red dots: returns to the location of the sugar drop. (B) Overlap in trajectories and distribution of return directions Search trajectories of *Drosophila* showed a significant higher degree of overlap in light compared to dark condition (light; *n* = 6; dark; *n* = 7; Mann-Whitney U test *P* = 0.02; Dunn post hoc test dmin 3: *P* = 0.022; dmin 5: *P* = 0.035). Orco mutants tested under light conditions showed the tendency to have a higher degree of overlap than wild type flies tested in the dark and a lower degree of overlap than wild types flies tested in light. Search trajectories of honey bees showed a low degree of overlap and did not differ under different light conditions. Yellow: light condition; black: dark condition; blue: *Drosophila* Orco mutant. Distribution of return angle did not find any evidence for a bias in return direction in flies and bees (Kolmogorov-Smirnov test; *P* > 0.05). In flies, we only found one run with a biased return angle (dark condition), and in honey bees two runs with a biased return angle (light condition). Radius of outer circles: *Drosophila*: 6 mm; *A. mellifera*: 11 mm.

Closer analysis of the trajectories revealed that flies and honey bees in fact returned to the location of the sugar drop during their search. Furthermore, all detected returns were highly precise in the sense that flies and bees always entered a circular area with a radius of one body length around the sugar drop (red lines, Fig. 3*A* for details see *Experimental Procedures*). A similar observation was reported for *Musca domestica* (14). Both findings suggest that the sugar-elicited local search involves more complex spatial orientation, e.g. memory of the food location, than just an increase in turning frequency (3,4).

### Vision, olfaction, and gustation are not necessary to return to the sugar drop location

As flies and bees often returned to the location of the sugar drop, we asked which sensory channels they use to find back. One idea is that both species use chemosensory signals or cues, e.g. footprint pheromones or fragments of sugar-water (15-18). However, it is also possible that they use self-motion (idiothetic) information (19-24). As a first step to answer this question we tested whether the search trajectories showed any evidence of overlapping trails, which would be present if flies and honey bees marked their walking path. We calculated the amount of overlapping sections for single trajectories (line width = 1 pixel; *Drosophila* body width: ∼5 pixels; *A. mellifera* worker body width: ∼7 pixels) using two respectively three different trail distances (3 pixels ∼ 50% body overlap, 5 pixels = partial overlap, 10 pixels = no overlap; Fig. 3*B*). Interestingly, *Drosophila*, search trajectories under light condition showed a significant higher degree of overlap compared to trajectories under dark condition at a line width of 3 pixel (light: *n* = 6; dark: *n* = 7; Mann-Whitney U test *P* = 0.02; Dunn post hoc test: *P* = 0.022; Fig. 3*B*). Similarly, trajectories of the olfactory blind Orco mutant under light conditions showed a trend to have a higher degree of overlap than trajectories in the dark. Both results suggest that flies predominantly use vision instead of olfaction if they follow their own footprints. In honey bees, the trajectories showed a very low degree of overlap independent of the lighting condition (Fig. 3*B*).

For the analysis of overlap we omitted the close vicinity of the sugar drop because that region evidently showed a high degree of overlap. To test whether flies and bees might use chemosensory signals or cues close to the sugar drop location we examined if they used preferred directions to approach the sugar drop position. We tested whether in a single search run all the crossing points with a virtual circle around the sugar drop (*Drosophila*: 6 mm; *A. mellifera*: 11 mm) were randomly distributed (Fig. 3*B*). In flies as well as in bees the overall majority of individuals showed a random distribution of return angles irrespective of the lighting conditions (*P* > 0.05 Kolmogorov-Smirnov test; Fig. 3*B*).

Based on our results, we propose that sugar-elicited search behavior in flies and bees does not involve trail marking. Trail marking likely conflicts with the transitory duration of the local search as well as the goal to search in different directions (3). Furthermore, returns to the sugar drop location are hardly affected by close range chemosensory cues. However, it seems that *Drosophila* and to a lesser degree honey bees can and do follow their own trails using vision, but it appears that they do not systematically use them to find back to the starting point of the trajectory.

### Experimental removal of potential gustatory, olfactory and visual cues indicates that flies and bees use self-motion information to return to the location of the sugar drop

Finally, we performed experiments in which we presented the sugar drop on a removable band or a disc (*Drosophila*, band width = 5 mm; *Drosophila*: disc radius = 9 mm); *A. mellifera*: disc radius = 15 mm) to eliminate any chemosensory signal or cues on the ground of the arena. These experiments were either done in a homogeneous light environment (*Drosophila* band) or in the dark (*A. mellifera* and *Drosophila* disc experiments). In all experiments, in which flies and bees initiated a searching behavior and were motivated to return to the starting point (*Drosophila* band *n* = 7; total number of returns = 14; *A. mellifera*: *n* = 6, total number of returns = 12; Fig. 4*A*; see movie S7), we found that they returned to that location with the similar precision (Fig. 4*A*; see movie S8). Minimum distance between the sugar drop location and the head of flies or honey bees were not significantly different between light, dark and removable band or disc condition.

**Fig. 4.**
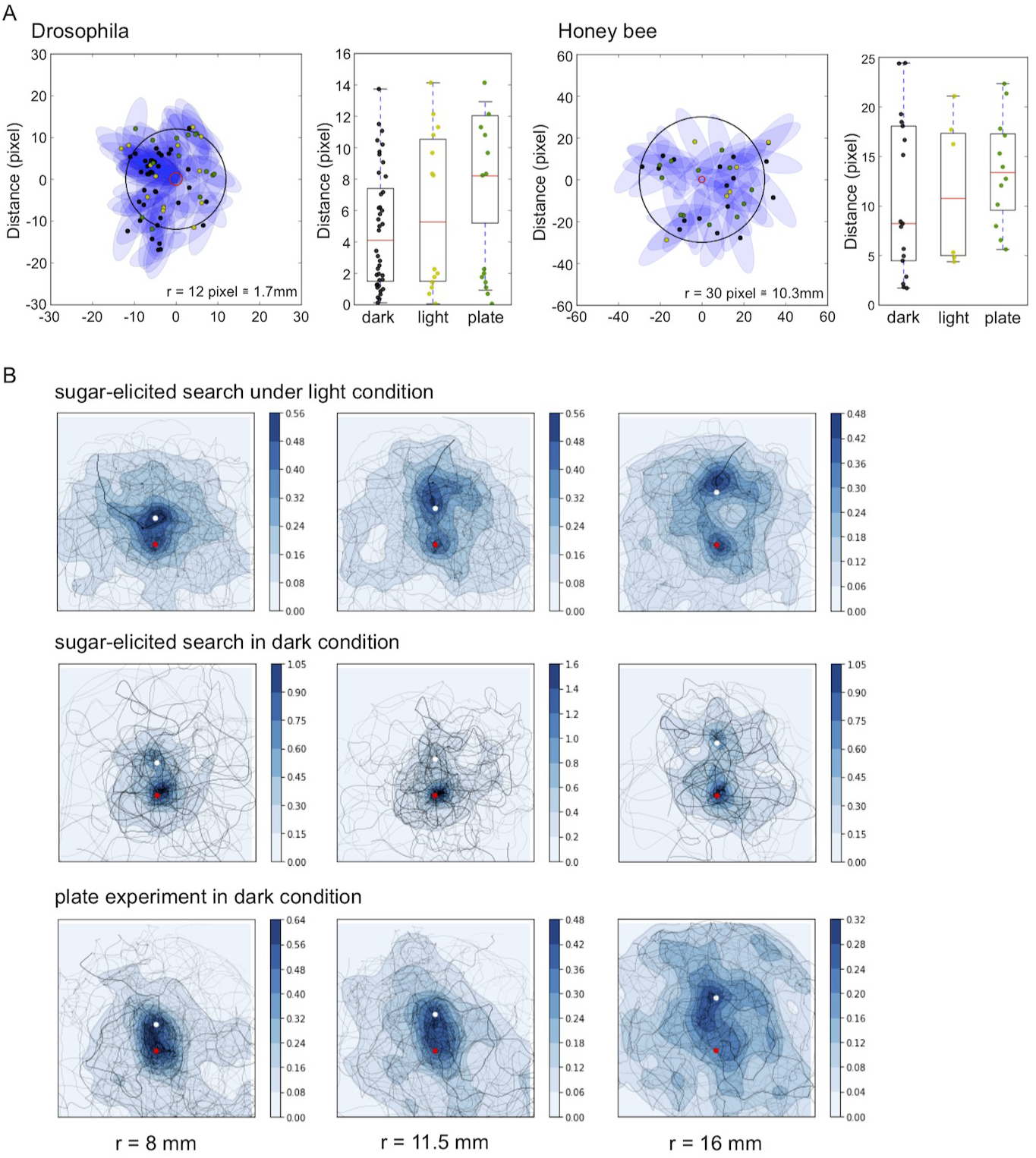
Removable band and disc experiments. (A) Comparison of the return accuracy under different experimental conditions: dark, light and removable band or disc. For Drosophila as well as honey bees, minimum return distances to the starting position did not differ between returns made under light, dark or the removable plate condition. Black circles: *Drosophila* r = 2 mm, *A. mellifera* r = 10 mm; Colored dots: black = dark condition; yellow = light condition; green = stripe (*Drosophila*) or disc (*A. mellifera*) experiment. (B) Probability heatmaps of the spatial distribution of *Drosophila* flies after defining a new virtual starting point (light condition *n* = 20; dark condition *n* = 20; disc experiment *n* = 20). For each experiment we plotted heatmaps for three new starting positions defined by radial distances of 8 mm, 11.5 mm, and 16 mm from the original sugar drop location. Red dot = sugar drop location; white dot = new starting position.

There are two possible mechanisms how flies and bees return to the location of the food source: (a) the turning behavior increases the probability to return to the starting position of the search trajectory (25), or (b) flies and bees use self-motion (idiothetic) information and path integration to intentionally return to the location of the food source. To decide between these two hypotheses we compared the probability of returning to a randomly selected point of the trajectory with the probability of coming back to the original food location. We did this analysis for three Drosophila experiments (light *n* = 20, dark *n* = 20, and removable disc *n* = 20). The new fictive starting position of the search was defined as the point where the trajectory crossed a virtual circle with a selected radius (*Drosophila*: radius = 8 mm, 11.5 mm, or 16 mm; Fig. 4*B*). Then, we rotated the trajectories so that the new starting point was mapped into a predefined point and superimposed all runs to generate a heat map. If returning to the location of the sugar drop was the result of a higher turning frequency (i.e. change in walking pattern), then the original sugar drop location should be indistinguishable from the mean return probability to any other points equally distant from the new starting point. A higher than average return probability would indicate that returning to the location of the sugar drop can not be explained by a higher turning frequency and would indicate that flies and bees use path integration to return to the sugar location. For all Drosophila experiments we found a significant higher visitation probability for the location of the sugar drop than other equivalent points (Mann-Whitney U and t-test; dark and light, max *P* = 0.008, with disk max *P* = 0.04). For bees we did not see a strong statistical difference. As mention above honey bees constantly moved around and thus showed a relatively high visitation probability of visit for any given location in the arena.

Together, the results of our different experimental approaches provide evidence that flies and honey bees use self motion information (i.e. path integration) during sugar-elicited local search behavior (19-25). In this case the location of the sugar-drop likely functions as a reference point to organize a meaningful search around this location (26).

### Responses to visual cues and landmarks indicating the location of the sugar drop

Even if flies and bees do not need visual cues or landmarks to return to the food source, still they might be used and modulate the sugar-elicited search trajectory. To test the effect of visual cues on the search behavior, we first presented flies and bees with a black dot beneath the sugar drop. For flies, the presentation of a single black dot did not affect the SeS index (no dot: *n* = 31 median SeS = 0.46 ± 0.30, black dot: *n* = 39 median SeS = 0.40 ± 0.33). The number of individuals initiating returns was slightly but not significantly increased (no dot: 12.9%, black dot: 23.07%). Interestingly, the distribution of returns was bimodal and flies that had experienced a dot underneath the sugar drop performed a higher number of late returns (no dot: number of late return = 1, black dot: number of late return = 7, *P* = 0.03 Mann-Whitney U test). These late returns might be due to an innate attraction for visual contrast or a learnt response towards a black dot similar to that one they had encountered during food intake (26). The strongest effect of the black dot was a significant reduction in the mean radial distance of the trajectories, indicating that the flies spend more time in the vicinity of the sugar location (no dot: median 13mm ± 8 mm; black dot: median 5.5 ± 4 mm; Mann-Whitney U test *P* = 0.013; Fig. 5*A*, see movie S9).

**Fig. 5.**
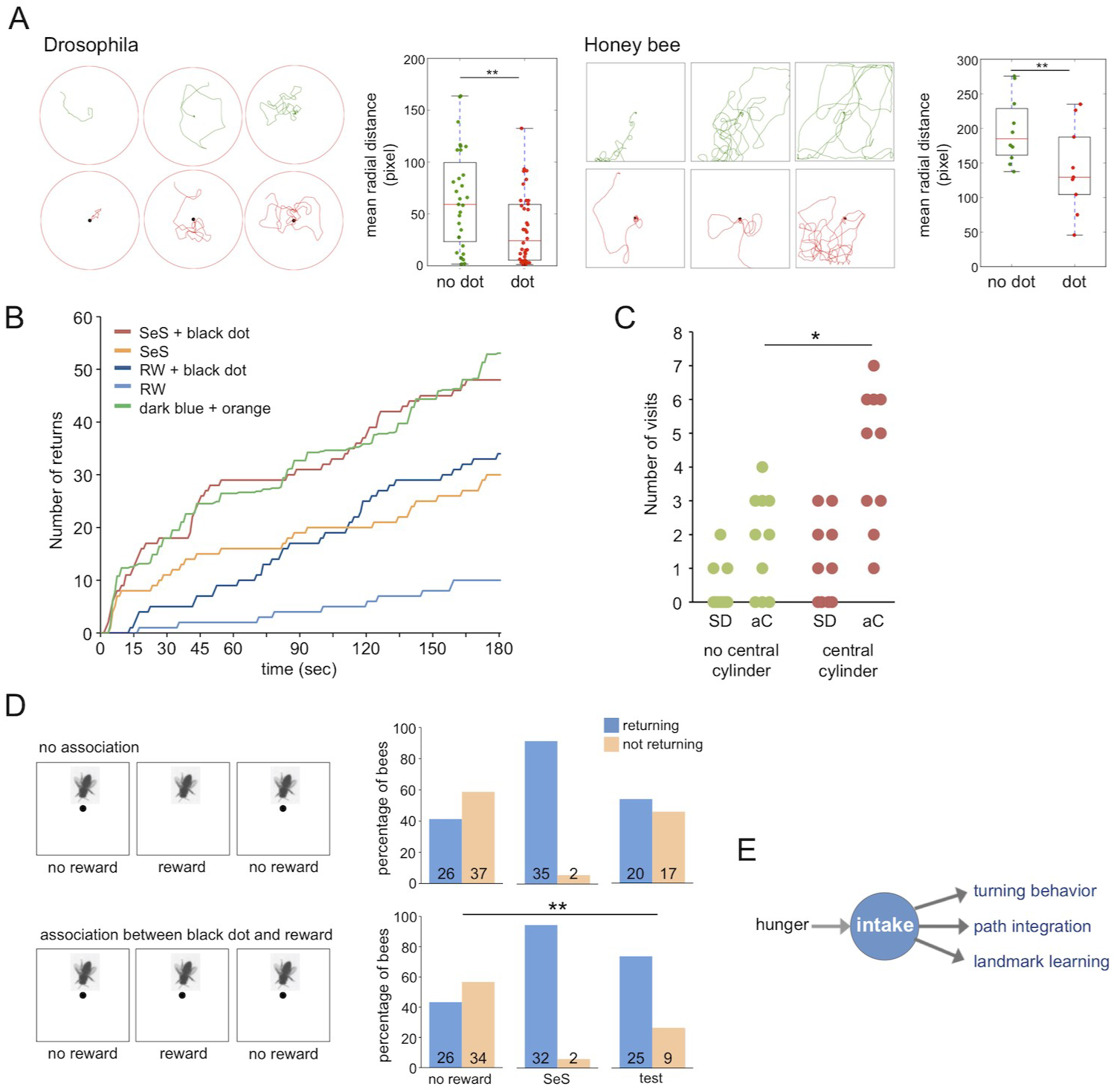
Responses to visual cues and landmarks indicating the sugar drop location. (A) Black dot beneath the sugar drop led to a reduction in the mean radial distance in flies no dot: median 13mm ± 8 mm; black dot: median 5.5 ± 4 mm; Mann–Whitney U test *P* = 0.013; and bees (no dot: median 63 ± 11 mm; black dot: 44 ± 18 mm; Mann–Whitney U test *P*_(2)_ = 0.045). (B) Cumulative return number to an unmarked or visually marked location in random walk and sugar-elicited search behavior. Light blue: random walk without black dot (*n* = 10); dark blue: random walk with black dot (*n* = 10); orange: sugar-elicited search behavior without black dot (*n* = 10); red: sugar-elicited search behavior with black dot (*n* = 10); green: best fit linear combination of random walk with dot and SeS without dot. (C) Number of visits to peripherally located black cylinders after honey bee workers encountered or did not encountered a similar black cylinder in the vicinity of the sugar drop. Green: experiment without a cylinder next to the sugar drop (*n* = 10); Red: experiment with a a cylinder next to the sugar drop (*n* = 10; student t-test *P*_(2)_ = 0.004). SD: sugar drop; aC: additional cylinders. (D) Associative learning experiment. Individual bees were tested in three consecutive trials (3min, intertrial interval = 3 min). Trial 1: spontaneous return response towards an unrewarded black dot. Spontaneously responding bees were excluded from trial 2 and 3. Trial 2: Presentation of a sugar drop and induction of sugar-elicited search. Control group: no black dot (*n* = 37); test group: black dot underneath the sugar drop (*n* = 34). Trial 3: Test run, presentation of an unrewarded black dot. The associative learning trial changed the proportion of individuals responding to the black dot without any reward (2-sample Z test one-tailed, Z-Score = -2.8236, *P* < 0.005). In addition, the response probability is also significantly different from the response probability after sugar stimulation alone (Z-Score one-tailed= -1.7016. *P* = 0.045). (E) Scheme: Behavioral responses involved in sugar-elicited search.

Similar to *Drosophila*, honey bees did not show a significant change in the search behavior in the presence of a single dot (no dot: *n* = 10 median SeS = 0.49 ± 0.12, black dot: *n* = 9 median SeS = 0.52 ± 0.15). In contrast to flies, the number of individuals initiating returns was equally high in both conditions (no dot: 78%, black dot: 80%). The distribution of returns during the search was slightly bimodal and numbers of late returns were slightly higher in the presence of a black dot. Similar to *Drosophila*, the black dot led to a significant reduction in the mean radial distance of the trajectories in *A. mellifera* (no dot: median 63 ± 11 mm; black dot: 44 ± 18 mm; Mann–Whitney U test *P* = 0.045, Fig. 5*A*, see movie S10).

To explore the effects on visual cues or landmarks in greater detail we performed three additional experiments with honey bees using a closed arena, which allowed studying behavioral responses for a longer time (3 min). In the first experiment we compared the responses of workers to a black dot during random walk and sugar-elicited search behavior. A black dot in the center of the arena was highly attractive for honey bees exploring the arena without a sugar stimulation (RW compared with RW + dot, Mann– Whitney U test *P* < 0.0001). Further, a black dot did also increase the number of returns in the sugar-elicited search assay (SeS compared with SeS + dot, Mann–Whitney U test *P* < 0.0001). However the differences in the return frequency induced by a black dot did not differ between random walk and sugar-elicited search (Fig 5*B*). This finding suggest that the intake of the sugar drop does not significantly increase visual attention in this behavioral context.

Then we asked whether the experience of a black cylinder (height: 15 mm; diameter: 6 mm; see photo S13) close to the sugar drop affect the number of visits to similar additional cylinders in the arena. Honey bees that encountered a black cylinder next to the sugar drop visited significantly more often distant landmarks than those that did not encounter a landmark (no central cylinder; *n* = 10; central cylinder; *n* = 10; Student’s *t* test *P*_(2)_ = 0.004; Fig. 5*C*, see movie S11).

Finally, we asked more specifically whether the intake of the sugar drop is capable of inducing associative learning. Two groups of bees were exposed to three consecutive experimental runs (intertrial interval 3 min; Fig. 5*D*): (1.) a spontaneous response trial, in which bees were released over a black dot but without a sugar reward, (2.) a learning trial, in which one group of bees were presented with the black dot and a drop of sugar water and a control group of bees with a drop of sugar water but no black dot, and (3.) a test trial, in which the bees were again presented with the black dot but no reward. If honey bees learn to associate the black dot with a sugar reward in the second trial, a higher number of bees that received the black dot pairing should initiate a return behavior in the third experimental run. 43% (respectively 41%) of the naïve foragers showed at least one spontaneous “return” to the black dot in the first run, and in the second run 94% (respectively 95%) of the bees, that did not show a spontaneous return in the first run, initiated a sugar-elicited search behavior with returns. After the learning trial more than 74% of the tested bees responded to the black dot in the third run. In the non-associative experiment 54% of the bees responded to the black dot in the third run. The distribution of “returning” and “non-returning” bees in the test run is significantly different from that in the spontaneous run (2-sample Z test two-tailed, Z-Score is -2.8236, *P* < 0.005).

Based on the results of these diverse experiments, we conclude that hungry bees, and likely also flies, already show a heightened visual attention towards conspicuous visual cues when exploring an area. The intake of a drop of sugar does not significantly increase this visual attention. However, the intake of sugar is capable to induce an association learning of nearby landmarks that then affects the search trajectory. In this context it is interesting to note that Fukushi (27, 28) already used the sugar-elicited search assay to test color learning in the blow fly *Lucilla cuprina*. To summarize, sugar intake in the sugar-elicited search behavior initiates not only path integration but also landmark learning.

## Discussion

The significant finding of our experiments is that sugar-elicited search behavior, first demonstrated by Vincent Dethier, is way more complex than previously proposed and comprises a set of complementary behavioral responses including increase in turning frequency, path integration and landmark learning (Fig. 5*E*). Thus, we show that this local search behavior involves two behavioral strategies that play a major role in large-scale insect navigation (1,5,7,8). For the future, sugar-elicited search assay provides the opportunity to use *Drosophila* and its neurogenetic toolkit to study the neural circuits and genetic mechanisms underlying food search behavior, navigation, and path integration (12, 29-31).

White et al. (14) were the first to suggest that sugar-elicited search behavior involves active returning to the location of the sugar drop. Reasons to return to the original location might be the probability that the food source is not depleted or has been replenished. Flies and bees likely use several different sensory systems, e.g. vision, olfaction, and gustation, to find back to the food location. However, our removable plate experiments under dark conditions clearly demonstrate that they are capable of finding back in the absence of visual, olfactory, and gustatory cues. Thus, flies and honey bees also use self-motion (idiothetic) cues, e.g. proprioreceptor input, to navigate back to the sugar location in the dark. The most basic definition of path integration is keeping track of one’s own movement using self generated (idiothetic) motion signals to be able to return back to the starting point of that movement irrespective of the distances travelled (19-21). Recently, Zeil and colleagues (22) provided some evidence that the Banded Sugar Ant (*Camponotus consobrinus*) likely uses path integration during local search behavior supporting our finding that path integration can be used during small-scale search behaviors. Probably any kind of locomotion is supported by a short-term memory of idiothetic information as a record of previous movements necessary for guiding current and future movements (3) During evolution, insect central place foragers might have elaborated this system to be able to use path integration mechanisms to navigate over larger distances (7,8).

Interestingly, there is some evidence that honey bees are capable of using walking path information to generate dance information (32,33). Thus, the neural circuits involved in sugar-elicited search behavior, walking path integration, flight navigation, honey bee dance, might be connected or even be the same (34). In this case, sugar-elicited search assay provides the opportunity to use *Drosophila* and its neurogenetic toolkit to study the neural circuits and genetic mechanisms underlying food search behavior, navigation, and path integration (12,29-31). Parallel experiments in honey bees will allow to determine the behavioral differences between flies and a master of insect navigation, as well as verify whether the behavioral responses in the lab assay correspond to those used in large scale navigation in nature (35-37).

## Materials and Methods

### Drosophila melanogaster

Male flies of the Canton-S strain of *Drosophila melanogaster* were used throughout. Flies were raised on glucose-cornmeal-yeast-wheat germ medium at 25°C, and a 12 hour light/dark cycle. Behavioral experiments were done during the central 4 hours of the light period. Flies eclosed within 12 hours periods were collected and maintained in a fresh medium for one day and afterwards starved for 24 hours in a vial with a water (evian®) soaked Kimwipe paper at the bottom.

For the experiments, single flies were transferred into 0.5 ml micro centrifuge tubes. The tube was placed on the LED lighting box for more than 3 minutes for acclimatization. A Petri-dish was placed on the lighting box and a tiny amount of silicone oil was placed in the center of the Petri-dish to hold the 0.1 µl of 200 mM sucrose solution. The solution was colored with a blue food dye to detect the presence of sugar solution. Then we placed the tube over the sugar droplet and waited until the fly found the droplet. Immediately after the fly started to ingest the solution, we removed the tube and surrounded the Petri-dish with a white cylindrical tube (67 mm inner diameter × 100 mm height, polyvinylchloride resin) and started the video recording (Logicool HD Webcam C615, or NET COWboy DC-NCR20U, Digital Cowboy). Recordings were done for 3 min at 30 f/sec.

Experiments under dark conditions were performed in a dark room (< 1 lux, measured by CENTER 337 digital mini luxmeter; Center Technology Corp., Taiwan) and the arena was illuminated with infrared LEDs (850 nm, S8100-60-B/C-IR, Scene Electronics, China). Spectrometer (QE65000, Ocean Optics, U.S.A.) measurements confirmed that the spectrum ranged from about 730 nm to 930 nm with a maximum emission at 846 nm. Preparation and positioning of the flies were done under the dim deep red light. When the fly commenced ingestion of the solution and we turned off the light and started the video recording (DC-NCR20U, Digital Cowboy or Flea3, Point Grey (40 f/sec, 1214 mm lens, Azure).

We did two sets of experiments to demonstrate that flies are capable of returning to the sugar drop location without visual and chemosensory cues. The first set of experiments was done using a homogeneous light condition (Petri-dish surrounded with a white cylindrical tube, 67 mm inner diameter × 100 mm height, polyvinylchloride resin). The sugar drop was presented on a removable band (band width 5mm). The second set was done in dark conditions (see above) and the sugar drop was presented on a transparent disc (small disc: 5.6 mm diameter, large disc: 17 mm diameter). The discs were 0.175 mm in thickness and made of clear polyester sheet. The disc was removed when the fly started walking and left the disc.

For the landmark experiments we used a blue rectangular cellophane film (2 x 2 mm) which was taped to the bottom of the Petri dish.

All experiments belonging to one set were over several days, but the respective test conditions were alternated.

### Honey bees (A. mellifera)

Experiments were done using nectar foragers of *A. mellifera* colonies that were kept on the NCBS campus, Bangalore. Colonies were provided with pollen and sugar-water (1 M sucrose) at artificial feeders. Foragers were caught with centrifuge tubes when landing on the feeder before they started to collect sugar-water. Then they were brought to the laboratory and placed in tube on the arena. After three minutes we started the experiment. The sugar drop (3 ul, 2 M sucrose solution) was positioned in the centre of the arena (31.5 x 31.5 cm). When the bee started feeding, the tube was removed and the video recording started (40 f/sec, Flea3, Point Grey, 1214 mm lens, Azure). Experiments under dark conditions were done in a dark room (< 1 lux, measured by CENTER 337 digital mini luxmeter; Center Technology Corp., Taiwan) and the arena was illuminated with infrared LEDs (850 nm, S8100-60-B/C-IR, Scene Electronics, China). Spectrometer (QE65000, Ocean Optics, U.S.A.) measurements confirmed that the spectrum ranged from about 730 nm to 930 nm with a maximum emission at 846 nm. Spectral sensitivity of the long-wavelength green receptors in *A. mellifera* workers ranges from 330 - 650 nm with a maximum at 544 nm (38,39). In the learning experiments we used landmarks composed of a black dot (diameter: 10 mm) and a vertical black cylinders (height: 15 mm; diameter: 6 mm). All experiments stopped when the bees left the arena. After each run the arena was cleaned with 70% ethanol.

### Analyses of trajectories

Walking trajectories for flies and honey bees were generated using ctrax software (40; http://ctrax.sourceforge.net/). We conceived a new set of MATLAB and Python routines for the analyses of the sugar-elicited search trajectories.

#### 1. *Sugar-elicited search index* (SES-index)

We developed an index of convolution as a parameter of the intensity of search behavior. The formula of the the SeS-index is defined as a function of the time average:

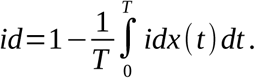

The position of the insect at time t is (x(t),y(t)). The position is calculated from the sugar drop location, i.e. the head position of the fly or honey bee while feeding. The distance of the insect from the center is given by 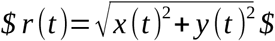 and total length of the trajectory traversed by the insect is given by d(t). We defined the SeS-index as the average of a geometric ratio 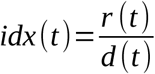. The SeS-index will give a value between 0 and 1 as r(t)<d(t). For an exact straight line trajectory the SeS-index = 0; for an insect moving in a sufficiently convoluted trajectory, the SeS-index would approach 1. We numerically calculate d(t) and then numerically integrate the SeS-index ratio to get SeS-index. The calculation of the SeS-index starts after the fly or honey bee moved away one fly/bee size from the sugar drop location. As *Drosophila* showed a lot of longer stops of grooming, we also excluded these parts from the calculation of the SeS-index.

#### 2. Identification of returns to the sugar-drop location

For the algorithm identifying returns in the search trajectories, we defined two concentric circles. An inner circle indicating the position of the sugar drop (radius r0; *Drosophila* r0 = 2.5 mm; A. mellifera r0 = 7 mm), and the outer circle indicating the minimum distance that a fly or honey bee had to move away from the sugar drop location (radius r1; *Drosophila*: r1 = 4 mm; *A. mellifera*: r1 = 12 mm). A return is a movement out of the outer circle (r1) and then coming back into the inner circle (r0).

#### 3. Distribution of return directions in the vicinity of the sugar drop location

For a given trajectory, we note all it’s crossings with the inner and outer circle and see if the angles of these crossing are non-randomly distributed.

#### 4. Identification of overlapping sections in the search trajectories

Ctrax generated trajectories have a width of a single pixel (the center of the fly or bee) and they hardly overlap. We take into account the actual size of a Drosophila male (width: 5 - 6 pixels) and an A. mellifera worker (width: 10 - 12 pixels) and consider a selected trajectory width (3, 5, or 7 pixels) to calculate the overlap. As the overlap is meant as a proxy for trail following, we also defined a minimum duration for an overlap (= 0.5 sec, i.e. 15 frames for flies, and 20 frames for honey bees).

## Acknowledgements

The idea to study the sugar-elicited search behavior in honey bees goes back to discussions between AB and Gene E. Robinson (UIUC). AB, JJH and Jonathan Massey developed the set-up for honey bee experiments in Gene E. Robinson’s lab. We thank William McFadden, and Saud Sadulkar for getting started with Ctrax and the analyses of trajectories. Abhishek Anand helped with the honey bee experiments. The research project was supported by National Center for Biological Sciences (TIFR) institutional funding (to AB), by International Centre for Theoretical Studies (TIFR) institutional funding (to PB), and a Grant-in-Aid for Scientific Research by the Ministry of Education, Culture, Sports, Science and Technology of Japan to TT.

## Supporting Information

Movies

S1 Drosophila: water stimulated

S2 Drosophila: sugar stimulated

S3 Drosophila: in the dark

S4 Drosophila: removable sheet

S5 Drosophila: black dot

S6 Honey bee: sugar stimulated

S7 Honey bee: in the dark

S8 Honey bee: removable disc

S9 Honey bee: black dot

S10 Honey bee: landmarks without central landmark

S11 Honey bee: landmarks with central landmark

S12 Photos of the areana and landmarks used in the learning experiment

